# Relating Ecological Diversity to Genetic Discontinuity across Bacterial Species

**DOI:** 10.1101/2023.09.29.560152

**Authors:** Hemanoel Passarelli-Araujo, Thiago M. Venancio, William P Hanage

## Abstract

Bacterial genetic discontinuity, representing abrupt breaks in genomic identity among species, is crucial for grasping microbial diversity and evolution. Advances in genomic sequencing have enhanced our ability to track and characterize genetic discontinuity in bacterial populations. However, exploring systematically the degree to which bacterial diversity exists as a continuum or is sorted into discrete and readily defined species remains a challenge in microbial ecology. Here, we aimed to quantify the genetic discontinuity (*δ*) and investigate how this metric is related to ecology. We harnessed a dataset comprising 210,129 genomes to systematically explore genetic discontinuity patterns across several distantly related species, finding clear breakpoints which varied depending on the taxa in question. By delving into pangenome characteristics, we uncovered a significant association between pangenome saturation and genetic discontinuity. Closed pangenomes were associated with more pronounced breaks, exemplified by *Mycobacterium tuberculosis*. Additionally, through a machine learning approach, we detected key features that impact genetic discontinuity prediction. Our study enhances the understanding of bacterial genetic patterns and their ecological implications, offering insights into species boundaries for prokaryotes.

## INTRODUCTION

Bacteria exhibit remarkable genetic makeup and ecological versatility, thriving in diverse niches worldwide. Plummeting sequencing costs have led to a wealth of genomic data, enabling genetic diversity and evolutionary relationships to be explored across many bacterial species. However, an essential inquiry in microbial ecology pertains to whether bacterial genetic diversity exists as a continuum or as distinct species groups^1, 2, 3^.

The definition of bacterial species faces challenges because bacteria can exchange genetic material through horizontal gene transfer (HGT)^4^, potentially blurring the species boundaries. This complexity has led to divergent views on bacterial species existence: while it was once thought that excessive recombination would preclude their species formation^5^, a contemporary perspective suggests that the gene flow patterns can even delineate species^6^. Besides, recent studies have revealed a clear genetic discontinuity across bacterial genomes, supporting the existence of discrete genetic clusters (which might or might not correspond to species)^7, 8, 9, 10^.

Genetic discontinuity refers to the occurrence of a significant difference in genetic makeup between populations or groups of organisms^11^, thereby signifying potential boundaries between distinct species. This discontinuity can occur over time through natural selection, genetic drift, or geographic isolation. Besides, genetic discontinuity can be an important factor in determining whether populations should be classified as separate species^8, 10^.

Defining bacterial species is important for reasons beyond any human desire to catalog bacterial diversity; it is vital for understanding how evolutionary forces shape genetic lineages^12, 13^. Furthermore, proper classification impacts practical applications in industry, agriculture, and medicine. For instance, *Gardnerella vaginalis* illustrates the clinical relevance of properly naming individual species. Formerly grouped under *G. vaginalis*, the division into multiple species has revealed diverse health associations^14^, including species linked to bacterial vaginosis as opposed to those found in healthy vaginal microbiomes^15^. Therefore, the previous classification of all *Gardnerella* as *G. vaginalis* limited the ability of clinicians to assess when and whether the presence of *Gardnerella* indicated a health risk.

One way to assess species boundaries is by estimating the genetic relatedness between genomes. A robust method to classify bacterial species is based on the Average Nucleotide Identity (ANI) estimate, with organisms belonging to the same species if the possess around 95% ANI or more among themselves^4, 5^. Since estimating ANI for thousands of genomes is computationally expensive, alternative methods were developed to accommodate the growing genomic dataset^10, 16^.

Despite observed breaks in genetic identity distributions for various species, quantifying the magnitude of these breaks and their ecological implications remains a challenge. Here, we address the intricate nature of bacterial diversity through an empirical genomic distance network approach and pangenome analysis. We aimed to quantify the extent of genetic discontinuity within and between bacterial populations to determine whether intrinsic genetic boundaries can provide a more ecologically relevant basis for species redefinition. Here, we seek not to pigeonhole bacterial species, but to examine the presence of genetic boundaries, quantify their extent, and explore their ecological implications for species classification.

## Methods

### Data collection and network analysis

We download a dataset comprising 210,129 genomes available on RefSeq as of September 2022. To ensure data quality, we retained genomes with fewer than 500 scaffolds and utilized the GTDB quality control^17^ to exclude genomes displaying low quality or contamination. Filtered genomes were used to perform tens of billions of comparisons with mash v2.2.2 ^16^ to construct a weighted network (*M*) with igraph v1.5.1 ^18^. *M* comprises genomic relationships among the set of genomes (*g*), with edges (*e*) corresponding to the genomic identity between pairs of genomes, quantified as the inverse of Mash distance (1 - Mash).

The weighted network was used to select representative genomes to infer the genetic identity patters. To select representative genomes, we subsetted the network *M* to create *M*′:

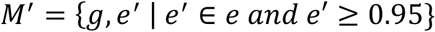

Therefore, *M*′ represents a species network containing the same set of genomes, but retaining edges above 95% identity, a threshold used to define species^4^. We employed the label propagation algorithm^19^ to detect communities in *M*′ that represent species. Only communities containing more than 50 genomes were used. This criterion was adopted to mitigate downstream modeling errors and enhance confidence in the biological significance of species representation.

Representative genomes of each species were chosen based on the following criteria: (i) prior designation as representative in both GTDB and RefSeq databases, (ii) representative at least in GTDB, or (iii) fewest scaffolds if the previous criteria were not met. Ties were resolved by random selection. This yielded a subset of 261 representative genomes (*t*) used for subsequent analysis.

### Genome annotation, pangenome, and phylogenetic analysis

Genomes assigned to communities containing representative genomes were annotated using Bakta v1.5.1^20^. To ensure consistency in annotation, all genomes were reannotated within the same framework, mitigating potential discrepancies. In cases where communities exceeded 500 genomes, we implemented a downsampling strategy. Genomes showing 99.5% or higher identity were removed. For communities still surpassing this threshold, the remaining genomes were randomly selected.

We employed Panaroo v1.2.10^21^ in the moderate mode to obtain the pangenome. The R packages Pagoo^22^ and Micropan^23^ were used to estimate the pangenome openness and genomic fluidity, respectively. We used Orthofinder v2.5.4^24^ to obtain the orthogroups from representative genome communities. All single-copy genes were with Mafft v7.505^25^ and concatenated to reconstruct the phylogenetic tree with IQ-Tree v2.1.4^26^, incorporating ModelFinder^27^ to identify the best fitting model. One thousand bootstrap replicates were generated to assess the significance of internal nodes. The phylogenetic tree was visualized with ggtree^28^.

### Genetic discontinuity estimation (*δ*)

To estimate and quantify the genetic discontinuity across species, we employed an egocentric-based approach using representative genomes as “baits” within the network *M*. For each genome (*i*) in the representative subset (*t*), *i* served as an egocentric node to calculate its genomic distance (*d*_*i*_(*g*)) from all other genomes *g* in *M*. These distances *d*_*i*_ (*g*) were sorted in descending order to retrieve genomes most similar to the representative genome *i*. The sorted array (*D*_*i*_) yielded a ranked list of genomic similarities. For each index *j* in *D*_*i*_, the corresponding genomic identity (*I*_*i*_ (*j*)) was determined, quantifying the similarity between the representative genome *i* and the genome at index *j* in the array.

To assess the Genetic Rate of Change, we calculated the first derivative of the genomic identity (*I*_*i*_ (*j*)) with respect to *j*. The first derivative Δ_*i*_ was calculated as the change in genomic identity between two consecutive indices *j* and *j* + 1 in *D*_*i*_, divided by index difference:

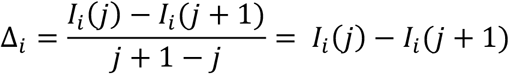

This rate of change Δ_*i*_ offered insights into how rapidly the genomic identity changed as we moved through the sorted array (*D*_*i*_), offering a measure of variability in genomic similarity.

Finally, let *δ* represent the genetic discontinuity, reflecting the idea of a break or sharp change in the genetic identity between a species and its closest relative. For each species, *δ* was defined as the maximum Genetic Rate of Change above 94%. This threshold was adopted to exclude breaks representing higher taxonomical classifications, focusing solely on the species level. For instance, consider hypothetical genera G1 and G2: *δ* captures the steepest change in genomic identity from species in G1, not those from G1 to G2 (see Figure 1). Thus, *δ* can be calculated as:

**Figure 1.**
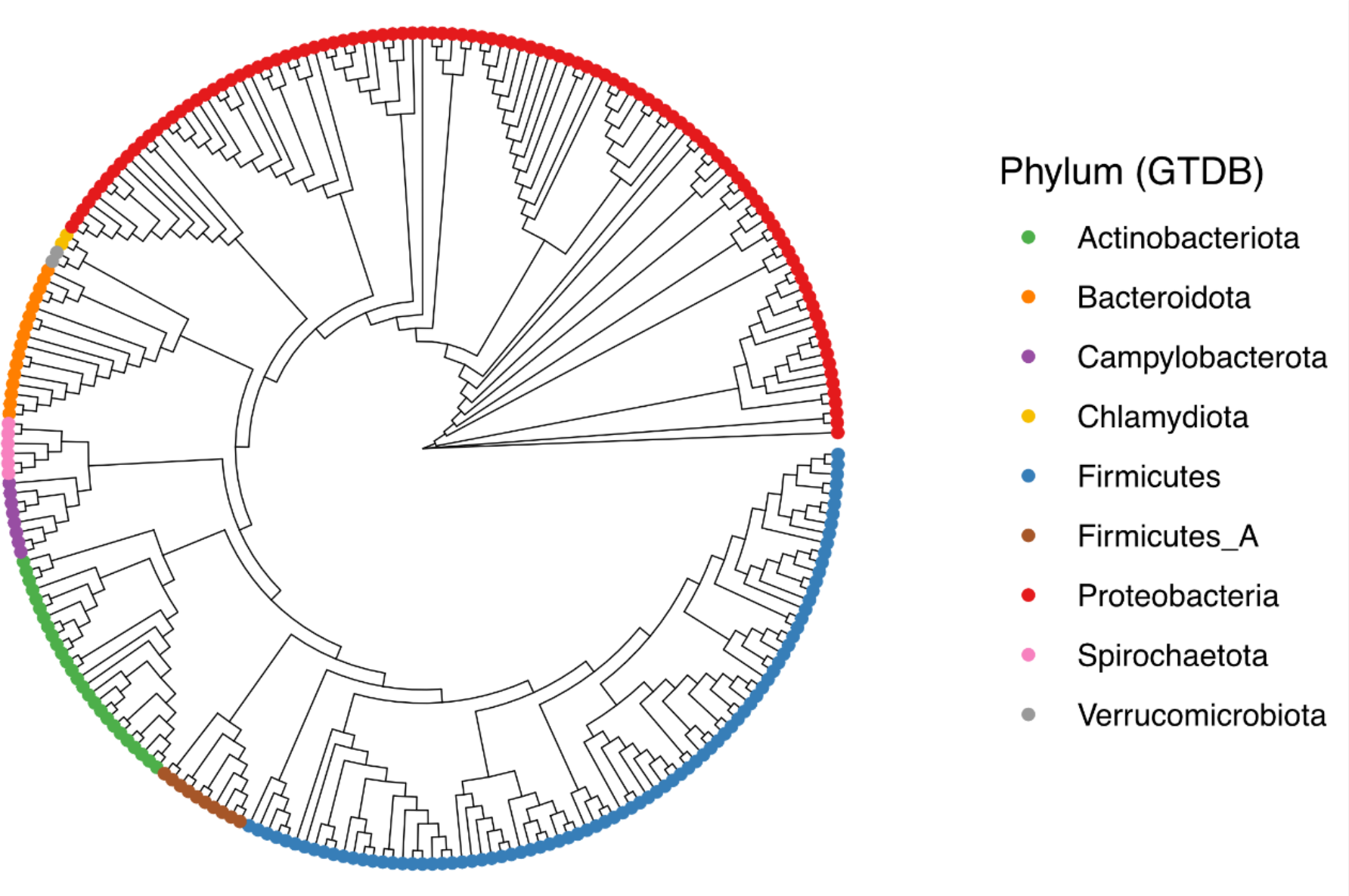
Phylogenetic tree of representative species. The phylogenetic tree reconstructed from single-copy genes of 261 representative species used in this work to explore bacterial genetic discontinuity.

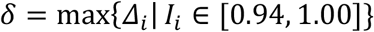

Additionally, we also incorporated GTDB species classification as a second prior to enhance accuracy in modeling *δ* for downstream analyses.

### Machine Learning modeling

The outcome prediction task was formulated as a regression problem. We tested four different ML models to predict the genetic discontinuity (*δ*) for each species: Linear Regression, Lasso Regression, Support Vector Regressor, Random Forest Regressor, and Gradient Booster Regressor. This analysis was employed using the Scikit-learn v1.0.2^29^ and XGBoost v1.5.2^30^ python^31^ libraries.

We considered 261 species and 31 features categorized into four main groups: taxonomical, orthology-related, annotation-based, and pangenome metrics. Taxonomical features encompassed GTDB classification for each species (phylum, class, order, family, and genus). Ortholog-related features refers to six metrics obtained after detecting orthogroups for reference genomes (e.g., proportion of species-specific orthogroups) (Table S1). Categorical features with more than two categories were represented by a set of dummy variables, with one variable for each category.

Annotation-based features were retrieved from all 45,550 reannotated genomes. Continuous variables, including %GC content and coding density, were represented by their median value to account for their asymmetrical probability distribution. For discrete variables such as the number of CRISPR arrays, we calculated their relative frequency, indicating the likelihood of a species carrying such elements. This yielded 13 annotation-based features. Seven pangenome metrics, encompassing pangenome openness, genomic fluidity, and core genome proportion (core genes to pangenome size ratio), were included in the feature dataset.

Given the relatively low number of observations, the entire dataset was employed for training the model. We utilized the Extra Tree Regression feature selection method to reduce dimensionality, improve the estimator’s accuracy, and boost the model performance. This algorithm employs randomized feature selection and ensemble averaging to make predictions, helping identify influential features, reduce overfitting, and enhance the model’s performance^32^. Also, we adopted k-fold cross-validation to mitigate dataset size limitations and to evaluate model performance metrics (Root Mean Squared Error (RMSE), Mean Absolute Error (MAE), and Quantile Loss).

To tune hyperparameters in the final ML model, we conducted a grid search with k-fold cross-validation utilizing 5 folds. We used the RMSE as the model score metric. We retrieved the importance of variables on explaining the model, by adopting the SHAP (*Shapley Additive exPlanations*) technique. the essence of SHAP is to measure the feature contribution of each individual to the outcome and whether the feature has a positive or negative impact on predictions^33^.

## Results

### Dataset information

We obtained 258,603 genomes from RefSeq that were filtered following the GTDB protocol^17^. After removing low quality and fragmented genomes, we retained of 210,129 genomes to explore bacterial genetic discontinuity. According to the NCBI classification, the top four abundant species in our dataset are *Escherichia coli* (n = 22,853), *Staphylococcus aureus* (n = 12,747), *Klebsiella pneumoniae* (n = 10,387), and *Salmonella enterica* (n = 9,755).

Over 44 billion comparisons were performed to construct an identity matrix *M*. The next step was defining communities in this network. Representative genomes for each community were selected by removing edges below 95% identity in *M*, resulting in 7,122 communities identified using label propagation – a proxy for species number (see methods for more details). Notably, 84.84% of communities contained fewer than 10 genomes, consistent with prior observations in the genus *Pseudomonas*^34^. For instance, a previous study found that 29% of officially recognized *Pseudomonas* type strains appeared as isolated nodes in a similar network analysis, highlighting the substantial underestimation of diversity when relying solely on type strains^34^. By focusing on communities with over 50 genomes (3.85%), we obtained a set of 261 representative genomes, enabling meaningful comparative genomic analyses subsequentially.

We reannotated a total of 45,550 genomes, representing nine phyla according to GTDB classification (Figure 1). Out of 942,094 genes detected, 95.1% were successfully assigned across 39,552 orthogroups. Moreover, species-specific orthogroups represented approximately 0.9% of the genes. Single-copy genes were used to reconstruct the phylogenetic tree. Except for *Proteobacteria*, the tree indicated monophyly in all other phyla based on GTDB classification, contrasting with the polyphyly observed based on NCBI classification, as noted before^17^. The GC content in these genomes ranged from 25.9% in *Mesomycoplasma hyorhinis* to 73.4% in *Streptomyces albidoflavus* (Table S1). Notably, *Clostridioides difficile* exhibited the greatest number of CRISPR arrays (median = 8) in a community of 500 genomes, with at least one array each. The dataset included genomes with diverse sizes, from the smallest commensal *Metamycoplasma hominis* (0.7 Mb) to the larger free-living bacterium *Burkholderia cepacia* (8.5 Mb).

### Clear genetic discontinuity revealed across bacterial species

To explore genetic discontinuity across different species, we employed a strategy in which a representative genome serves as a “bait” node within the network and the identity of all other genomes from it is calculated. This method allowed us to assess the ranked identity distribution from each representative genome (Figure 2a), revealing clear breakpoints within the distribution. For instance, considering the representative species of *Acinetobacter baumannii*, the 4968^th^ genome maintains a 97.27% identity. However, the identity of the 4969^th^ genome drops drastically to 93.34%, exemplifying the observed genetic discontinuity or genetic break.

**Figure 2.**
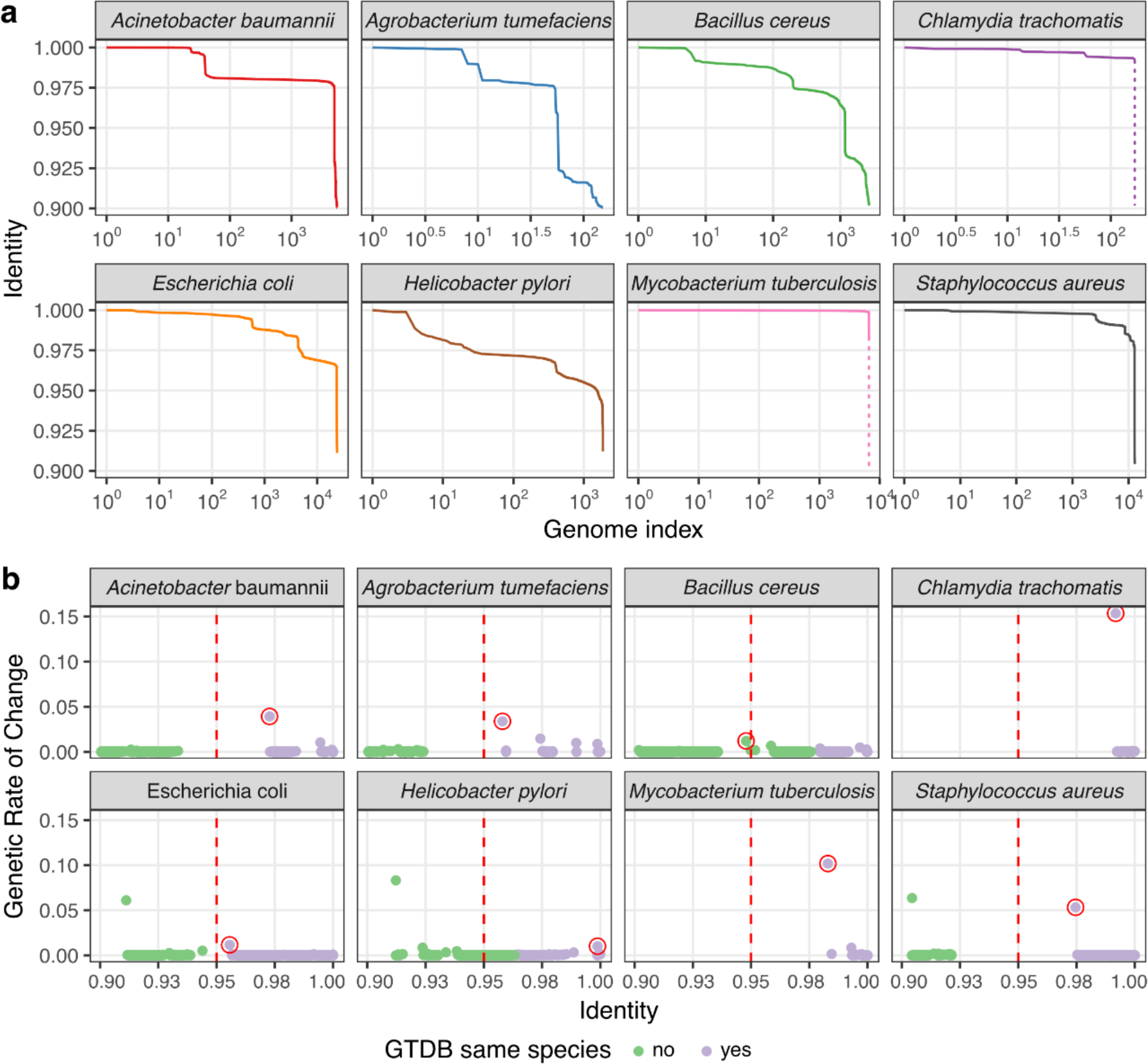
Genetic discontinuity properties of ten selected species. **a)** distribution of ranked genomic identities, revealing breakpoints around 95%. Vertical dashed lines for Chlamydia trachomatis and Mycobacterium tuberculosis indicate breaks beyond 90% from their closest genomes. The x-axis is represented in log-scale. **b)** Genetic Rate of Change is depicted across different identity values, with the break-associated point emphasized by a red circle. This point represents the key measure of genetic discontinuity (*δ*) examined in this work (see methods for comprehensive information). Genomes are colored based on GTDB classification.

We systematically quantified the genetic discontinuity by estimating how rapidly the genomic identity decayed as we moved through the sorted identity array. We took the first derivative of the distribution, offering a measure of variability in genomic similarity that we named as Genetic Rate of Change (GRC) (Figure 2b). The maximum value of GCR resulted in the genetic discontinuity metric *δ*, which characterizes the steepest change in genomic identity (see methods). For instance, the genetic break observed from 97.27% to 93.34% in *A. baumannii*, corresponds to *δ* = 0.9727 − 0.9334 = 0.0393 for this species (Table 1, Figure 2b).

**Table 1:** Genomic and ecological characteristics of 15 representative species. Genetic discontinuity (*δ*) represents the steepest change (break) in genomic identity distribution from a representative genome. The saturation coefficient (*α*) indicates pangenome openness: the higher the *α* value, the more closed the pangenome is. Core proportion refers to the ratio between the number of core genes and pangenome size of each species.

| Species | Lifestyle | $\delta$ | $\alpha$ | Core Prop. |
| --- | --- | --- | --- | --- |
| *Coxiella burnetii* | Obligate intracellular | 0.291 | 0.930 | 0.672 |
| *Treponema pallidum* | Obligate pathogen | 0.256 | 0.959 | 0.855 |
| *Mycoplasma pneumoniae* | Obligate intracellular | 0.228 | 0.967 | 0.755 |
| *Metamycoplasma hominis* | Obligate intracellular | 0.175 | 0.832 | 0.471 |
| *Chlamydia trachomatis* | Obligate intracellular | 0.154 | 0.976 | 0.895 |
| *Mycobacterium tuberculosis* | Obligate pathogen | 0.102 | 0.932 | 0.639 |
| *Staphylococcus aureus* | Opportunistic pathogen | 0.053 | 0.844 | 0.306 |
| *Acinetobacter baumannii* | Opportunistic pathogen | 0.039 | 0.073 | 0.152 |
| *Agrobacterium tumefaciens* | Plant pathogen | 0.034 | 0.764 | 0.312 |
| *Pseudomonas syringae* | Plant pathogen | 0.030 | 0.708 | 0.189 |
| *Rhizobium leguminosarum* | Symbiont | 0.024 | 0.704 | 0.230 |
| *Klebsiella pneumoniae* | Opportunistic pathogen | 0.016 | 0.730 | 0.161 |
| *Bacillus cereus* | Free-living | 0.012 | 0.640 | 0.085 |
| *Escherichia coli* | Free-living | 0.011 | 0.696 | 0.149 |
| *Helicobacter pylori* | Free-living | 0.010 | 0.725 | 0.189 |

Eight species were selected to showcase bacterial discontinuity, encompassing both pathogenic and non-pathogenic strains across various phyla and lifestyles (Figure 2b). Notably, *Chlamydia trachomatis* and *M. tuberculosis* exhibited pronounced *δ*, indicating substantial shifts in genetic similarity. Conversely, *Helicobacter pylori* represented few instances where a species lacks a clear genetic discontinuity, suggesting a blurred genetic boundary possibly influenced by its evolutionary history and lifestyle.

### Higher genetic discontinuity associates with allopatric lifestyle

To explore the ecological implications of bacterial discontinuity, we analyzed the pangenome of the 261 species mentioned above, which provides valuable insights into their lifestyles and evolution^35, 36^. The pangenome encompasses the core genome (genes present in all isolates), the accessory genome (genes in more than one but not all isolates), and isolate-specific genes. Pangenome openness, measured by the saturation coefficient (*α*), indicates the extent to which new gene families are detected in the pangenome as more genomes are included. Higher *α* values suggest gene pool saturation (the addition of new genomes contributes fewer new detected genes), while lower *α* values imply a more flexible genomic repertoire.

We used the pangenome openness to indirectly assess the species lifestyle^35^ (Table 1). A low saturation coefficient (open pangenome) suggests a flexible genomic repertoire, characteristic of sympatric populations that frequently exchange genes to adapt to various environments. Conversely, a high saturation coefficient (closed pangenomes) is associated with allopatric populations adapted to specific niches with limited gene exchange due to physical isolation or genetic incompatibility^35, 36^.

In our investigation, we identified a noteworthy correlation between pangenome openness and genomic fluidity (*ϕ*) (Figure 3a) – a measure of genomic dissimilarity at the gene level ^37^. Specifically, we found a negative correlation, indicating that species with closed pangenomes exhibit lower genomic fluidity, as previously noted^38^. Furthermore, we observed a pronounced increase in genetic discontinuity as the pangenome saturation coefficient rises.

**Figure 3.**
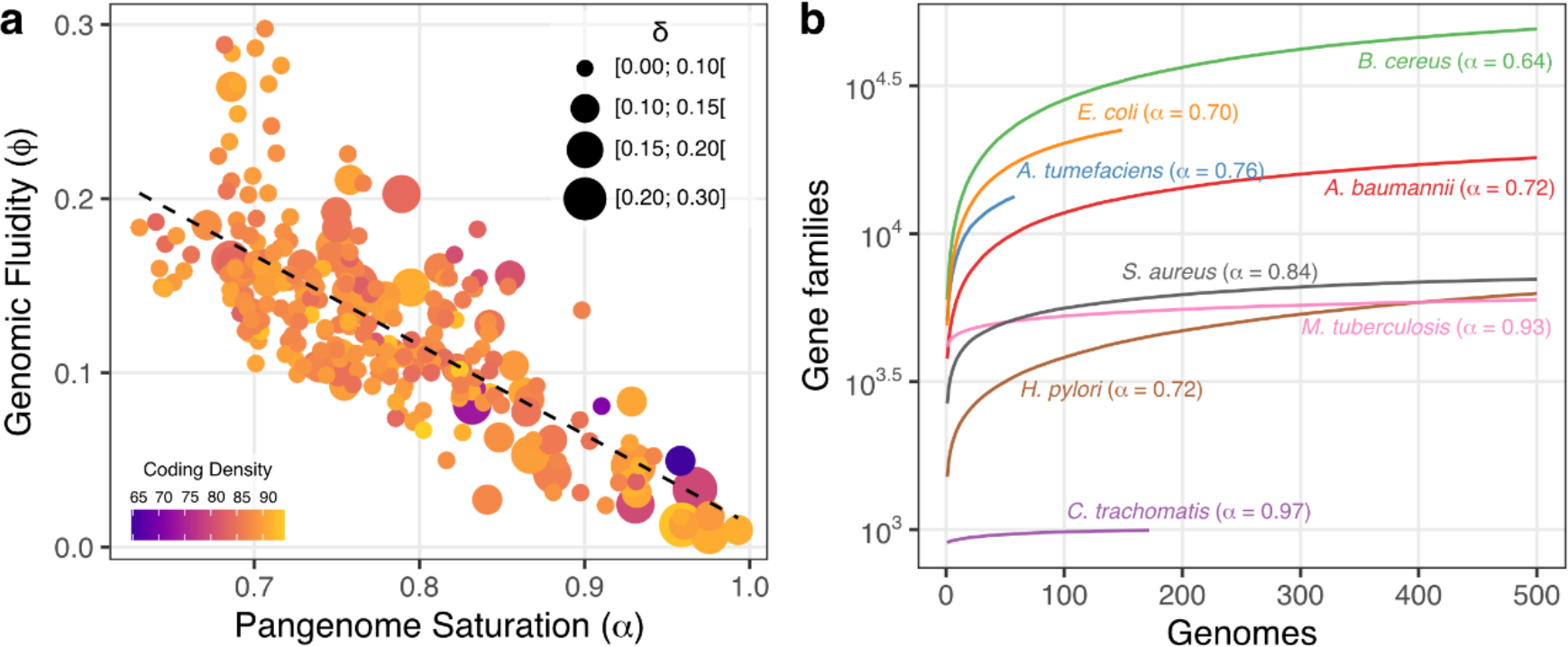
Pangenome properties of representative species. **a)** genomic fluidity in function of pangenome saturation with a linear regression line given by *ϕ* = −0.51148 × *α* + 0.52509. The size of the dots corresponds to different levels of genetic discontinuity, grouped into four categories. Additionally, the color of the dots indicates the coding density of each representative genome. **b)** pangenome openness of ten selected species, with *α* highlighted for reference.

*C. trachomatis* and *Bacillus cereus* exhibit distinct pangenome characteristics that reflect their contrasting lifestyles. *C. trachomatis* exhibited a closed pangenome (*α* = 0.97), indicating a limited capacity for gene acquisition through HGT. This suggests a relatively stable genome and a more specialized lifestyle, features associated with an obligate intracellular pathogenic behavior^39^. This pattern is frequent among species with high genomic discontinuity, closed pangenomes, allopatric lifestyles, and highly conserved pangenomes (Table 1). In contrast, *B. cereus* displays an open pangenome (*α* = 0.64), indicating a high propensity for gene acquisition and genomic diversity. This suggests a more versatile lifestyle, potentially enabling *B. cereus* to occupy various ecological niches and adapt to changing environments. The variations in pangenome openness observed in these two species provide valuable insights into their lifestyles and on how their pangenomes evolve in the context of genetic discontinuity.

### Uncovering most influential features to predict bacterial genetic discontinuity

We aimed evaluate the importance of different features in predicting bacterial genetic discontinuity. To address the asymmetrical nature of *δ* distribution (Figure 4a) and the impact of the number of genomes to estimate the pangenome openness, we employed a quantile regression that allowed us to assess the influence of pangenome saturation on genetic discontinuity while controlling for the number of genomes used to compute *α*. We found a significant impact across all quantiles examined (Figure 4b). Notably, as the quantile increased, the impact of pangenome saturation in *δ* became more pronounced. For instance, in the top quantile (τ = 0.95; [0.1040 − 0.291]), an increase of one unit in alpha corresponded to a 0.44-unit rise in *δ*. These results shed light on the pivotal role of pangenome saturation in shaping genetic discontinuity patterns.

**Figure 4.**
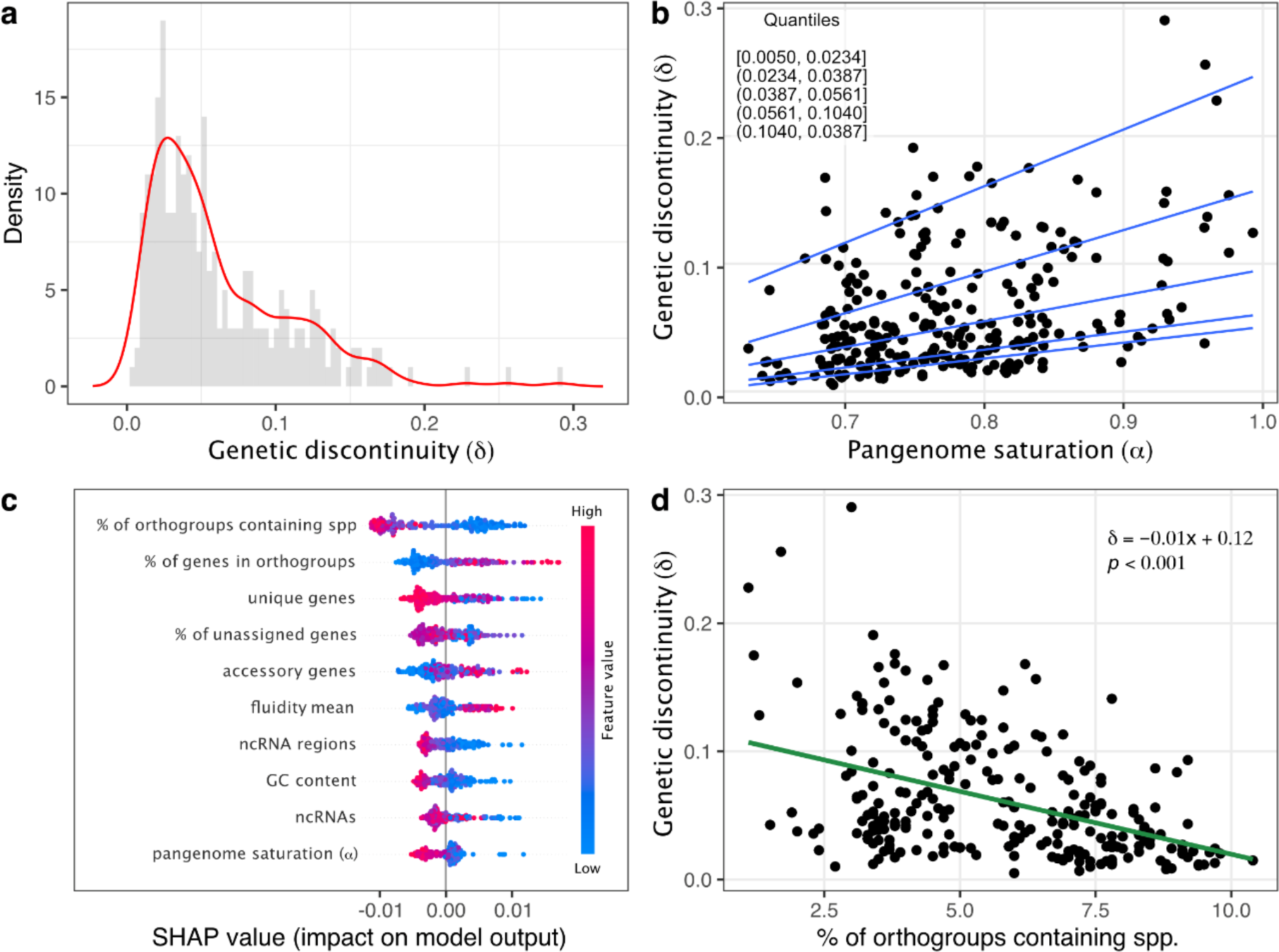
Genetic discontinuity modeling. **a)** Probability distribution of genetic discontinuity (*δ*) estimated from representative genomes. **b)** quantile regression analysis of *δ* as a function of pangenome saturation (*α*), highlighting a positive impact of *α* on *δ* across different quantiles (0.15, 0.25, 0.50, 0.75, 0.95). **c)** SHAP values derived from Random Forest Regression model, indicating the importance of ten features in predicting *δ*. Each representative genome is displayed, along with the positive or negative impact of each feature. **d)** linear regression between *δ* and the percentage of orthogroups containing species, the feature with the highest impact on predicting *δ*. The equation of the regression line and the associated p-value are also shown.

Beyond pangenome features, we also used a rich set of features ranging from taxonomical classification to orthology assessment to model bacterial discontinuity through a machine learning approach (see methods). Among the six tested methods, Linear Regression and Random Forest demonstrated superior performance in terms of root mean squared error, mean absolute error and quantile loss over different quantiles (Figure S1). Random Forest was chosen due to its ability to handle data distribution and collinearity without imposing strict assumptions. After hyperparameter tuning via k-fold cross-validation and grid search, we delved into feature importance using SHAP values to predict *δ*.

The most significant variable affecting the prediction of genetic discontinuity was the “Percentage of Orthogroups Containing Species” (Figure 4c). This metric gauges the proportion of orthogroups containing at least one gene from a given species. For example, if at least one gene from species A is present in 90 orthogroups from a total of 100, the percentage of orthogroups containing species A would be 90%. Moreover, this feature negatively impacts *δ* (Figure 4d; p-value <0.001). This metric helps associate the presence and representation of a species within orthogroups with genetic discontinuity, as it indicates how frequently genes from that species are involved in shared functional contexts across different organisms.

## Discussion

In this study, we systematically quantified genetic discontinuity (*δ*), which reflects significant shifts or abrupt changes in genetic similarity compared to a representative genome. Our work offers insights into the ecological roles of bacterial discontinuity and its implications for species classification. We carefully considered whether rigid taxonomic boundaries capture the fluidity and evolutionary dynamics that shape species’ histories, especially considering organisms where genetic exchange, adaptation, and hybridization prevail^40, 41^.

Philosophically, species classification raises the issue of Aristotelian essentialism – the idea that there are inherent qualities that define a species^42, 43^. The act of assigning species to specific categories becomes an exercise in grappling with the fundamental question of what defines a species. Is it solely genetic similarity, shared phenotypic traits, ecological niche, or something deeper that eludes our current understanding? In this work, we deployed genomics and ecology approaches to assess species boundaries. We used the essentialism idea as a prior to select representative genomes and explore whether the resulting genetic variation exhibited continuity or discreteness. To be agnostic about the choice of representative genomes, we employed a network approach where we could also retrieve genomes from understudied populations to represent a given community, avoiding the limitation of exploring species based only on well-studied type strains^34^.

Bacterial genetic discontinuity has already been observed in studies comprising thousands of genomes by using different approaches^7, 8, 34^. The remaining inquiry was how to measure genetic discontinuity and examine its ecological significance, while accounting for potential external factors such as sampling biases. To address this, we devised a novel metric (*δ*) by examining the maximum value in the first derivative distribution of genome identity derived from a representative genome. We observed delta values spanning from 0.005 (*Acinetobacter pittii*) to 0.290 (*Coxiella burnetii*) – species with remarkably distinct lifestyles. For the extreme case *C. burnetii*, the idea of *δ* = 0.29 means that the most similar genome in a different community within a network of over 200 thousand genomes shares only 29% genetic identity with the representative genome of the *C. burnetii* community, which comprises 65 genomes. This identity value is way beyond of what we expect to distinguish species and approaches to those used to define higher taxonomical ranks such as genera and families^44, 45^. *C. burnetti* is an obligate intracellular pathogen responsible for causing the Q fever disease in humans^46^. Its allopatric lifestyle may explain its high genetic discontinuity.

After identifying breaks that varied according species lifestyles (Table 1), the next challenge was to assign an ecological meaning to them. The pangenome analysis is essential to gain insights into lifestyle and evolution a species^35, 36^. A key result we found here was that the greater the bacterial discontinuity, the more closed the pangenome was, always controlling for the number of genomes used to estimate the saturation coefficient. Besides *C. burnetti, M. tuberculosis* and *C. trachomatis* also illustrate situations where the magnitude of the break is related to lifestyles, especially regarding the HGT dynamics.

Conversely, species with ubiquitous or environmental lifestyles such as *E. coli* and *B cereus*, presented smaller breaks, but still well-defined boundaries. Those bacteria are known for their ability to thrive in various environments, including soil, water, and plant surfaces. Their diverse ecological niches might lead to more continuous genetic variation, exhibiting less pronounced breaks. In contrast, *Helicobacter pylori* posed an intriguing challenge with regards to genetic discontinuity. Our analysis revealed either an ambiguous or non-existent genetic break in this species, rendering it difficult to draw a line to delineate its boundaries. The absence of a discernible genetic break in *H. pylori* may be connected to its extremely high rate of homologous recombination and emphasizes the need for a nuanced approach to species boundaries and genetic cohesion, and suggests the presence of unique evolutionary dynamics that warrants further investigation.

Beyond the use of pangenome features to predict bacterial discontinuity, orthogroups assessment may be vital to understand genetic discontinuity. By using both the SHAP values and ExtraTreess Regressor to retrieve feature importance, the “percentage of orthogroups containing species”, assigned by orthofinder, was the most important variable. This variable reveals insights into genetic interconnectedness and shared functions among bacterial species within an ecological niche. A higher percentage indicates shared traits due to the frequent co-occurrence, suggesting ecological overlap. Conversely, a lower percentage implies species specialization, indicating distinct ecological roles. For example, species with closed pangenomes such as *C. trachomatis* had genes present in only 2% of the total number of orthogroups.

In conclusion, our study highlights the significance of bacterial genetic discontinuity in understanding microbial diversity and evolution. We have shown that closed pangenomes and pronounced genetic breaks correspond to specific bacterial lifestyles, offering insights into microbial adaptation. Furthermore, our findings emphasize the role of orthogroups in characterizing genetic discontinuity and ecological dynamics within bacterial communities. This study contributes to a more nuanced understanding of bacterial diversity, emphasizing dynamic genetic relationships and the need to reevaluate traditional species classifications in microbial ecology.

## Acknowledgments

We thank the Fulbright Brasil Commission for funding HP-A studies at Harvard T.H. Chan School of Public Health in 2022 and 2023. TMV research group is funded by Fundação Carlos Chagas Filho de Amparo à Pesquisa do Estado do Rio de Janeiro (FAPERJ; grant E-26/201.117/2022), Coordenação de Aperfeiçoamento de Pessoal de Nível Superio(CAPES; Finance Code 001), and Conselho Nacional de Desenvolvimento Científico e Tecnológico (CNPq).

## Supplementary Material

**Figure S1.**
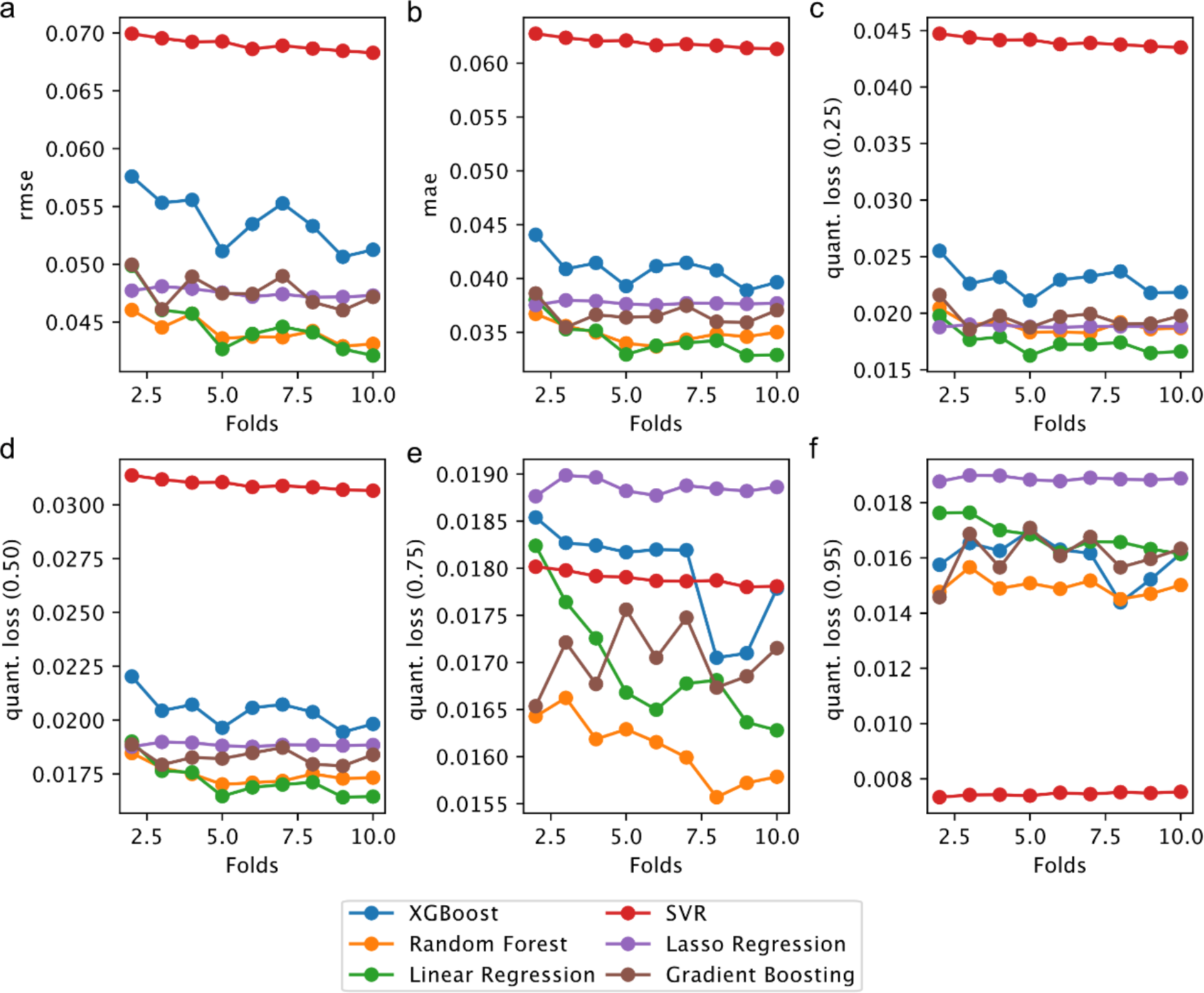
Model evaluation through k-fold cross validation using different metrics. **a)** Root Mean Squared Error (RMSE). **b)** Mean Absolute Error (MAE). **c-f)** Quantile Loss across four quantiles (τ = [0.25, 0.50, 0.75, 0.95]).

## References

1. Caro-Quintero, A. & Konstantinidis, K.T. Bacterial species may exist, metagenomics reveal. Environ Microbiol 14, 347–355 (2012).

2. Cohan, F.M. Systematics: The Cohesive Nature of Bacterial Species Taxa. Curr Biol 29, R169–R172 (2019).

3. Shapiro, B.J. et al. Population genomics of early events in the ecological differentiation of bacteria. Science 336, 48–51 (2012).

4. Bobay, L.M. The Prokaryotic Species Concept and Challenges. In: Tettelin, H. & Medini, D. (eds). The Pangenome: Diversity, Dynamics and Evolution of Genomes: Cham (CH), 2020, pp 21–49.

5. Doolittle, W.F. & Papke, R.T. Genomics and the bacterial species problem. Genome Biol 7, 116 (2006).

6. Arevalo, P., VanInsberghe, D., Elsherbini, J., Gore, J. & Polz, M.F. A Reverse Ecology Approach Based on a Biological Definition of Microbial Populations. Cell 178, 820–834 e814 (2019).

7. Knight, D.R. et al. Major genetic discontinuity and novel toxigenic species in Clostridioides difficile taxonomy. Elife 10 (2021).

8. Olm, M.R. et al. Consistent Metagenome-Derived Metrics Verify and Delineate Bacterial Species Boundaries. mSystems 5 (2020).

9. Passarelli-Araujo, H., Jacobs, S.H., Franco, G.R. & Venancio, T.M. Phylogenetic analysis and population structure of Pseudomonas alloputida. Genomics 113, 3762–3773 (2021).

10. Jain, C., Rodriguez, R.L., Phillippy, A.M., Konstantinidis, K.T. & Aluru, S. High throughput ANI analysis of 90K prokaryotic genomes reveals clear species boundaries. Nat Commun 9, 5114 (2018).

11. Hanage, W.P., Fraser, C. & Spratt, B.G. Sequences, sequence clusters and bacterial species. Philos Trans R Soc Lond B Biol Sci 361, 1917–1927 (2006).

12. Fraser, C., Alm, E.J., Polz, M.F., Spratt, B.G. & Hanage, W.P. The bacterial species challenge: making sense of genetic and ecological diversity. Science 323, 741–746 (2009).

13. Hugenholtz, P., Chuvochina, M., Oren, A., Parks, D.H. & Soo, R.M. Prokaryotic taxonomy and nomenclature in the age of big sequence data. ISME J 15, 1879–1892 (2021).

14. Potter, R.F., Burnham, C.D. & Dantas, G. In Silico Analysis of Gardnerella Genomospecies Detected in the Setting of Bacterial Vaginosis. Clin Chem 65, 1375–1387 (2019).

15. Hill, J.E., Albert, A.Y.K. & Group, V.R. Resolution and Cooccurrence Patterns of Gardnerella leopoldii, G. swidsinskii, G. piotii, and G. vaginalis within the Vaginal Microbiome. Infect Immun 87 (2019).

16. Ondov, B.D. et al. Mash: fast genome and metagenome distance estimation using MinHash. Genome Biol 17, 132 (2016).

17. Parks, D.H. et al. A standardized bacterial taxonomy based on genome phylogeny substantially revises the tree of life. Nat Biotechnol 36, 996–1004 (2018).

18. Csardi, G. & Nepusz, T. The igraph software package for complex network research. InterJournal Complex Systems, 1695 (2006).

19. Raghavan, U.N., Albert, R. & Kumara, S. Near linear time algorithm to detect community structures in large-scale networks. Phys Rev E Stat Nonlin Soft Matter Phys 76, 036106 (2007).

20. Schwengers, O. et al. Bakta: rapid and standardized annotation of bacterial genomes via alignment-free sequence identification. Microb Genom 7 (2021).

21. Tonkin-Hill, G. et al. Producing polished prokaryotic pangenomes with the Panaroo pipeline. Genome Biol 21, 180 (2020).

22. Ferres, I. & Iraola, G. An object-oriented framework for evolutionary pangenome analysis. Cell Rep Methods 1, 100085 (2021).

23. Snipen, L. & Liland, K.H. micropan: an R-package for microbial pan-genomics. BMC Bioinformatics 16, 79 (2015).

24. Emms, D.M. & Kelly, S. OrthoFinder: phylogenetic orthology inference for comparative genomics. Genome Biol 20, 238 (2019).

25. Katoh, K. & Standley, D.M. MAFFT multiple sequence alignment software version 7: improvements in performance and usability. Mol Biol Evol 30, 772–780 (2013).

26. Minh, B.Q. et al. IQ-TREE 2: New Models and Efficient Methods for Phylogenetic Inference in the Genomic Era. Mol Biol Evol 37, 1530–1534 (2020).

27. Kalyaanamoorthy, S., Minh, B.Q., Wong, T.K.F., von Haeseler, A. & Jermiin, L.S. ModelFinder: fast model selection for accurate phylogenetic estimates. Nat Methods 14, 587–589 (2017).

28. Yu, G. Using ggtree to Visualize Data on Tree-Like Structures. Curr Protoc Bioinformatics 69, e96 (2020).

29. Buitinck, L. et al. API design for machine learning software: experiences from the scikit-learn project. ECML PKDD Workshop: Languages for Data Mining and Machine Learning, 108–122 (2013).

30. Chen, T. & Guestrin, C. XGBoost: A Scalable Tree Boosting System. Proceedings of the 22nd ACM SIGKDD International Conference on Knowledge Discovery and Data Mining, 785–794 (2016).

31. Rossum, V. & Drake, F. Python 3 Reference Manual. Scotts Valley, California: CreateSpace; 2009.

32. Ferri, J., Pavel, P. & Hatef, M. Comparative Study of Techniques for Large-Scale Feature Selection. Pattern Recognition in Practice, IV: Multiple Paradigms, Comparative Studies and Hybrid Systems (2001).

33. Lundberg, S. & Lee, S. A Unified Approach to Interpreting Model Predictions. 31st Conference on Neural Information Processing Systems (2017).

34. Passarelli-Araujo, H., Franco, G.R. & Venancio, T.M. Network analysis of ten thousand genomes shed light on Pseudomonas diversity and classification. Microbiol Res 254, 126919 (2022).

35. Rouli, L., Merhej, V., Fournier, P.E. & Raoult, D. The bacterial pangenome as a new tool for analysing pathogenic bacteria. New Microbes New Infect 7, 72–85 (2015).

36. Brockhurst, M.A. et al. The Ecology and Evolution of Pangenomes. Curr Biol 29, R1094–R1103 (2019).

37. Kislyuk, A.O., Haegeman, B., Bergman, N.H. & Weitz, J.S. Genomic fluidity: an integrative view of gene diversity within microbial populations. BMC Genomics 12, 32 (2011).

38. Henaut-Jacobs, S., Passarelli-Araujo, H. & Venancio, T.M. Comparative genomics and phylogenomics of Campylobacter unveil potential novel species and provide insights into niche segregation. Mol Phylogenet Evol 184, 107786 (2023).

39. Stelzner, K., Vollmuth, N. & Rudel, T. Intracellular lifestyle of Chlamydia trachomatis and host-pathogen interactions. Nat Rev Microbiol 21, 448–462 (2023).

40. Arnold, B.J., Huang, I.T. & Hanage, W.P. Horizontal gene transfer and adaptive evolution in bacteria. Nat Rev Microbiol 20, 206–218 (2022).

41. Soucy, S.M., Huang, J. & Gogarten, J.P. Horizontal gene transfer: building the web of life. Nat Rev Genet 16, 472–482 (2015).

42. Austin, C. Aristotelian essentialism: essence in the age of evolution. Synthese 194, 2539–2556 (2017).

43. Shtulman, A. & Schulz, L. The relation between essentialist beliefs and evolutionary reasoning. Cogn Sci 32, 1049–1062 (2008).

44. Deloger, M., El Karoui, M. & Petit, M.A. A genomic distance based on MUM indicates discontinuity between most bacterial species and genera. J Bacteriol 191, 91–99 (2009).

45. Qin, Q.L. et al. A proposed genus boundary for the prokaryotes based on genomic insights. J Bacteriol 196, 2210–2215 (2014).

46. Seshadri, R. et al. Complete genome sequence of the Q-fever pathogen Coxiella burnetii. Proc Natl Acad Sci U S A 100, 5455–5460 (2003).

